# Biomechanics of shear-sensitive adhesion in climbing animals: peeling, pre-tension and sliding-induced changes in interface strength

**DOI:** 10.1101/031773

**Authors:** David Labonte, Walter Federle

## Abstract

Rapid control of adhesive forces is one of the key benchmarks where footpads of climbing animals outperform conventional adhesives, promising novel bio-inspired attachment systems. All climbing animals use shear forces to switch rapidly between firm attachment and easy detachment, but the detailed mechanisms underlying ‘shear-sensitive adhesion’ have remained unclear. Here, we show that attachment forces of stick insects follow classic peeling theory when shear forces are small, but strongly exceed predictions as soon as their pads start to slide due to high shear forces. Pad sliding dramatically increases the critical peel force *via* a combination of two distinct mechanisms. First, partial sliding will pre-stretch the pads, so that they are effectively stiffer upon detachment and peel increasingly like inextensible tape. We demonstrate how this effect can be directly related to peeling theories which account for frictional dissipation. Second, pad sliding reduces the thickness of the secretion layer in the contact zone, thereby decreasing the interfacial mobility, and increasing the stress levels required for peeling. The approximately linear increase of adhesion with friction results in a sharp increase of adhesion at peel angles less than ca. 30°, allowing rapid switching between attachment and detachment during locomotion. Our results may apply to diverse climbing animals independent of pad morphology and adhesive mechanism, and highlight that control of adhesion is not solely achieved by direction-dependence and morphological anisotropy, suggesting promising new routes for the development of bio-inspired adhesives.

---

Many insects, spiders, lizards and tree frogs can climb on plants and in the canopy of trees by employing adhesive footpads, which allow them to switch between strong attachment and effortless detachment within fractions of a second [1, 2, 3, 4]. The functional principles underlying this impressive dynamic control of attachment forces have attracted considerable interest amongst physicists, engineers and biologists, aiming to develop technical adhesives with similar properties [5]. A key feature of dynamic biological adhesive pads is that adhesive forces increase when they are pulled towards the body [6, 7, 8, 9, 10]. This simple, reversible and fast control mechanism which has been shown to have a much larger influence than retraction speed or normal pre-load [6, 8, 10]. Strikingly, shear-sensitive adhesion has been reported for ‘hairy’ and ‘smooth’ as well as ‘dry’ and ‘wet’ biological adhesive pads [6, 7, 8, 9], suggesting a universal control mechanism independent of pad morphology, and the alleged adhesive mechanism (van-der-Waals or capillary forces for dry and wet adhesives, respectively). What are the mechanisms giving rise to shear-sensitive adhesion?

Several previous studies have interpreted shear-sensitive adhesion of climbing animals using peeling theory (e. g. [6, 11, 12, 13, 14, 15, 7]), which predicts the force *F* required to peel off an elastic tape of width *w*, under a peeling angle *ϕ* (see inset in fig. 1 A). Assuming that the tape is infinitely flexible in bending, that deformations are in the limit of linear elasticity, and that effects due to inertia are negligible, the critical peel force per unit tape width, *P* = *F/w,* can be linked to the tape’s strain energy release rate *G* ([16, 17]; All equations are derived in detail in the Supplemental Material):

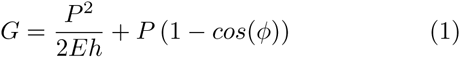

where *E* is the Young’s modulus of the tape, and *h* is its thickness. The first term on the right hand side, often called ‘elastic term’, is a combination of the energies (per unit area of detached tape) required to elastically stretch the detached fraction of the tape, and to move the point of force application due to tape stretching. The second term represents the work involved in moving the point of force application when a unit area of the tape is detached without stretching. For a thin tape of high stiffness, or for sufficiently large peel angles, eq. 1 approximately reduces to

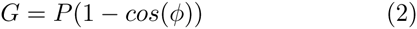

which we will refer to in the following as ‘inextensible tape model’, as it is exact for tapes of zero extensibility. Both eq. 1 and 2 predict that the peel force increases with shear force (as peeling occurs at smaller angles *ϕ*), but eq. 1 is limited by 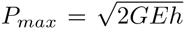 as *ϕ* → 0, while eq. 2 is unbound as a consequence of the assumption of infinite tape stiffness.

**Figure 1:**
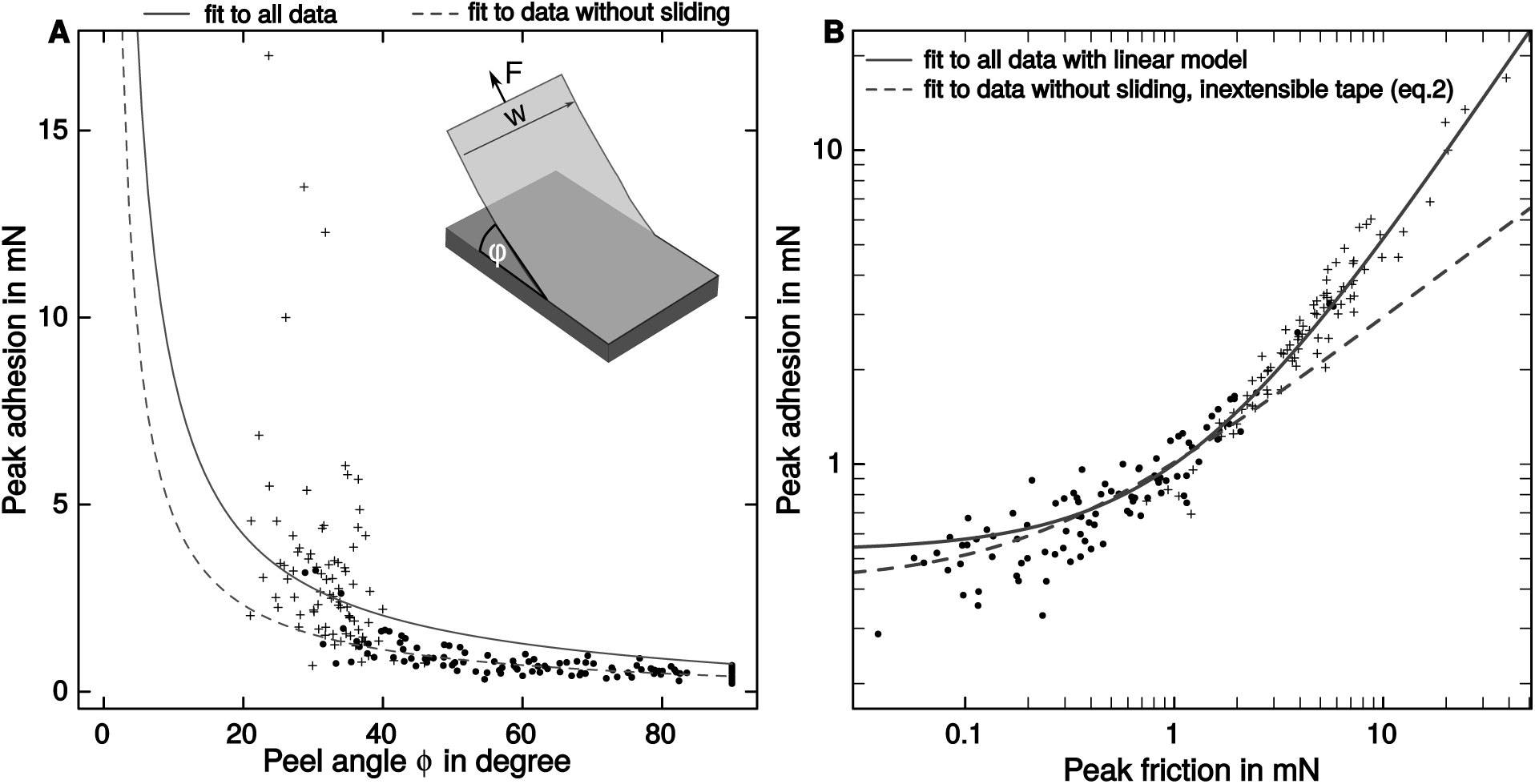
Individual pads of Indian stick insects were peeled off glass coverslips at different angles while measuring both adhesive and frictional forces. The symbols indicate whether peeling was accompanied by visible sliding (crosses for sliding or dots for static detachments, respectively). (A) Peak adhesion, *Fsin(ϕ),* against peel angle (n=11). The inextensible tape equation systematically overestimated adhesion for large peel angles (continuous line). For peel angles smaller than ≈35 °, most of the pads slid visibly during detachment. The model fit considerably improved when eq. 2 was restricted to data from measurements where no visible sliding occurred during detachment (dashed line). (B) Same data as in (A), but on a double logarithmic scale and with friction on the x-axis. The predictions of a simple linear model and the inextensible tape equation are similar for small shear forces (or large peel angles), but differ increasingly for large shear forces (or small peel angles). The divergence of the two models coincides with the onset of sliding.

Equation 1 has been used previously to study shear-sensitive adhesion in geckos, and tree frogs, but several problems arose [6, 7]. For example, the values for *G* and *E* required to fit experimental data exceeded plausible estimates, suggesting that additional dissipative mechanisms were at play [7]. In *Gekko gecko,* adhesion increased linearly with shear force, with a slope of around 0.5, indicating a constant ‘critical angle of detachment’ at *ca.* 30° (defined as the arc tangent of the ratio between adhesion and friction, see [6]). This is in contradiction to eqs. 1 and 2, as adhesive force, *F sin*(*ϕ*), cannot vary at a constant peel angle if *G* is constant. Autumn et al. [6] thus rejected tape peeling as an explanation for shear-sensitive adhesion in geckos. Several modifications of eq. 1 have been put forward since [11, 12, 13, 18], but the mechanics of shear-sensitive adhesion in insects, tree frogs and geckos still remain unclear.

Here, we study the biomechanics of controllable adhesion in stick insects *(Carausius morosus*). This article is organised as follows: First, we will show that shear-sensitive adhesion is consistent with peeling theory for large peel angles (or small shear forces), but is closer to a linear relationship between adhesion and friction for small peel angles (or large shear forces). Second, we will demonstrate that the departure from peeling theory coincides with the appearance of sliding during detachment, which sometimes led to re-attachment of previously detached parts of the adhesive pads. Third, we will use a simple first-principle modification of eq. 1 to discuss how ‘pre-strain’, sliding and ‘crack-healing’ can make even soft and thin tapes behave effectively as infinitely stiff. Lastly, we argue that this effect is still not sufficient to fully account for the discrepancy between peeling models and observed shear-sensitive adhesion. Instead, we provide evidence for a sliding-induced increase in interface strength, and suggest that in combination, the effects of sliding can account for the linear relationship between friction and adhesion observed in biological adhesives.

## Results&Discussion

The critical adhesive force, *F sin(ϕ),* required to peel off individual adhesive pads of stick insects from glass decreased significantly with the peel angle (fig. 1 A). A non-linear mixed model least squares fit of the inextensible tape model yielded a strain energy release rate of *G* = 1166mNm^−1^ (95% CI (1023,1309) mN m^−1^), not unusual for rubbery materials, but considerably higher than expected for vander-Waals forces (we justify the use of the inextensible tape equation below). This discrepancy is likely explained by viscous dissipation in the pad cuticle, as *G* approaches values typical for weak intermolecular forces in the limit of small peel velocities [10]. However, the inextensible tape fit systematically overestimated adhesion for larger angles, and underestimated forces for smaller angles (see fig. 1 A). In addition, we measured no peel angles (determined by the measured force vector) smaller than ≈ 22 °, despite two treatments which involved smaller surface ‘retraction angles’, indicating that some pads were sliding during detachment. Indeed, high-speed recordings of the contact area during detachment revealed that 81 out of 94 pads slid visibly when peeled at angles *ϕ* < 40 °. When the fit of eq. 2 was restricted to data from detachments without visible sliding (yielding *G* = 667 mN m^−1^, 95% CI (510,824) mN m^−1^), the agreement between theory and experiment considerably improved (fig. 1 A).

In contrast to the inextensible tape model, a simple linear model was in excellent agreement with the data, and explained around 95% of the overall variation in adhesion (see fig. 1B). A least-squares regression yielded a slope of 0.47, independent of whether pads were sliding during peeling (sliding vs. non-sliding, t_184_=0.74, p=0.46, n=11), and an intercept of 0.53mN (95% CI: (0.45,0.48) and (0.41,0.64) mN, respectively, fitted with a linear mixed model). Adhesion was approximately half of the acting shear force, indicating a critical detachment angle of ≈ 30 °, in remarkable agreement with earlier observations on gecko setae, despite the striking difference in pad morphology [19, 6]. However, we found significant adhesion in the absence of shear force (i.e. for *ϕ* = 90°, *t*_186_ = 8.98, p < 0.001, n=11, see fig. 1A), inconsistent with the phenomenological, zero-intercept ‘frictional adhesion’ model [6].

A plot of adhesion against friction on a log-log scale, along with a fit of (i) the inextensible tape equation restricted to detachment without sliding, and (ii) a linear model, shows that the predictions of both models are similar for small friction forces (or large peel angles; fig. 1 B). A comparison of the corresponding Akaike information criteria suggested that the inextensible tape model was in fact marginally better for friction forces smaller than approximately 2 mN (see Supplemental Material). For friction forces larger than approximately 2mN (or *ϕ* < 35 °), however, the model predictions differed increasingly, and the linear model was more accurate. The point of divergence coincided with the onset of sliding (see fig. 1 B).

### Pre-tension. partial sliding and ‘crack healing’

As the pads changed from static to dynamic contact, sliding of the entire pad was likely preceded by partial slippage close to the peel front. Such interfacial slippage can lead to a profound increase in the apparent strain energy release rate [20, 21, 22, 18, 23], as sliding ‘consumes’ part of the available energy, so that eqs. 1 and 2 are no longer valid. Gravish et al. [24] suggested that the adhesive strength of gecko setae is largely based on ‘external’ dissipation *via* seta sliding, superior to many commercial soft adhesives where interface toughness is largely based on ‘internal’ dissipation *via* plastic deformation, compromising structural integrity and thus limiting reusability. Indeed, the thin secretion layer covering the pads of all insects studied to date may serve as a lubricating ‘release layer’, helping to reduce viscous dissipation in the pad cuticle during voluntary detachment [10].

When a fraction of the attached pad slides, it will be stretched, resulting in an increase in the system’s elastic energy, and an associated movement of the point of force application. Remarkably, the energy loss by friction is as large as the corresponding change in the elastic term due to stretching ([5, 18] and see Supplemental Material). Upon detachment, the now pre-stretched pad extends less than an unstretched pad, and thus the work done by the applied load decreases. As a consequence, the required peel force increases - the interface gains strength. In this sense, peeling with frictional sliding is similar to the peeling of a tape which has been stretched *prior* to surface attachment, a case which has been thoroughly addressed by previous work [25, 26, 27, 28, 29, 13].

A quantitative assessment of the effect of pad pre-tension requires an approximation of the force that pre-strained the pad. For frictional sliding, this force is *F*_0_ = *F cos(ϕ)* (see Supplemental Material). In addition to frictional sliding, pre-tension may have arisen *via* one or a combination of two mechanisms in our experiments: First, low angle peeling can result in measurable strain in the pad cuticle [30]. Second, we observed ‘crack-healing’, i.e. previously detached parts of the pads reattached when peeling occured at low angles (see [25] for similar observations on rubber tapes, and fig. 2). In both cases, the peel force increased further after pre-tension was induced, so that *F*_0_ = *Fcos(ϕ)* is a plausible conservative estimate, independent of whether pads were stretched while in contact, or when detached. With this pre-strain eq. 1 becomes (see Supplemental Material)

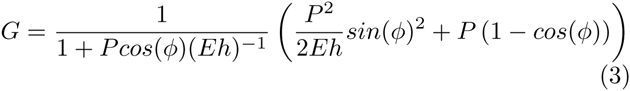

**Figure 2:**
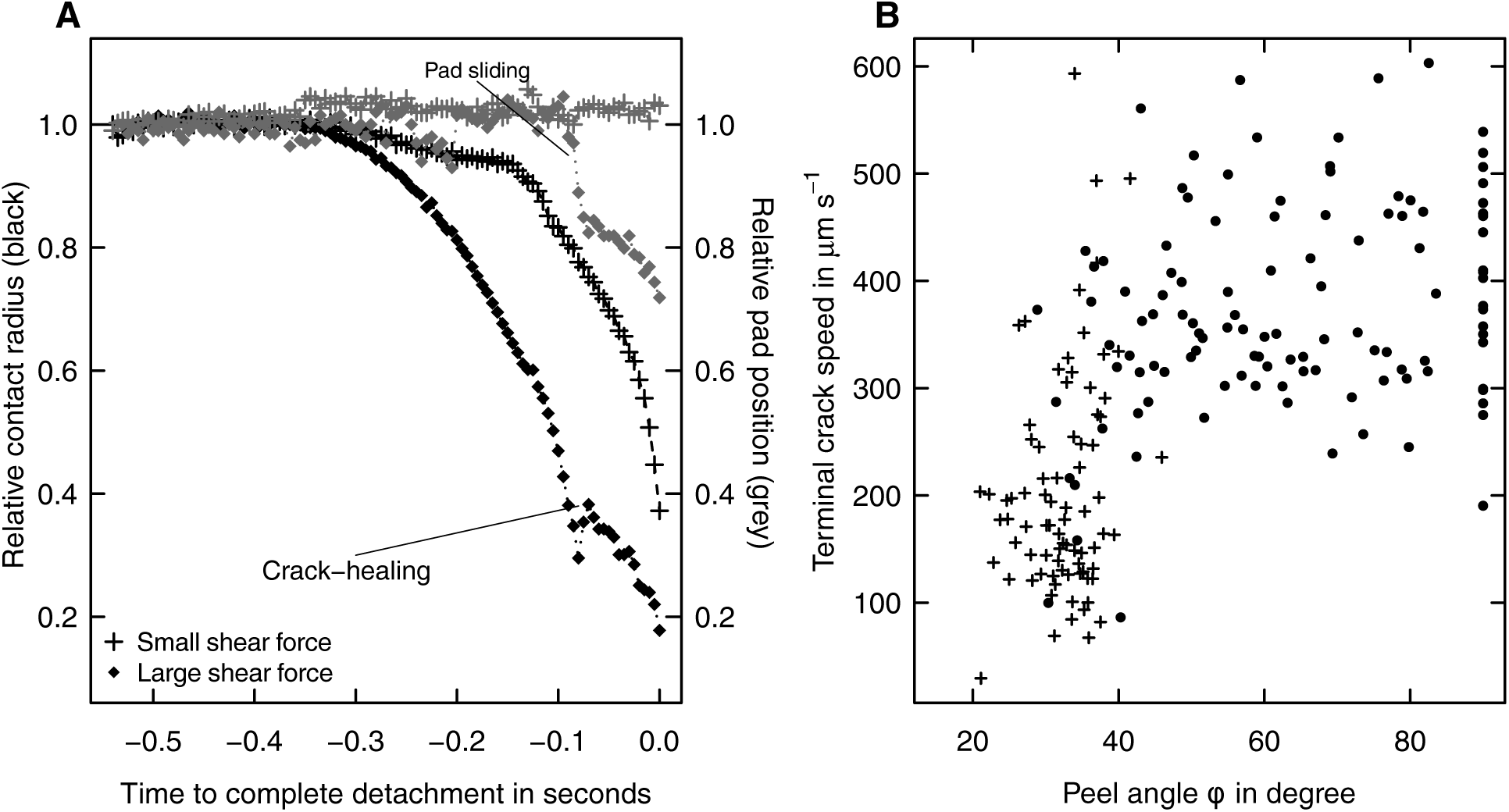
(A) During detachment, the contact radius of the pads – approximated as the ratio between pad area *A* and perimeter *Γ* – decreased continuously (black symbols show two example curves for shear forces of 0.5 and 4mN, respectively). The change of *A/Γ* with time may be interpreted as the speed of a crack propagating through the interface [31], which was measured by performing an ordinary least square regression of *A/Γ* against time for the last 50 ms of detachment (‘terminal’ crack speed). When shear forces were small, the crack initially accelerated, followed by approximately steady crack growth until detachment was complete. When shear forces were large, however, we sometimes observed that the crack was arrested or even receded, i. e. detached parts of the pad re-attached. This ‘crack-healing’ was clearly associated with the onset of sliding, i.e. the pad’s position relative to the surface changed (grey symbols). (B) As a result, detachments with visible sliding (crosses) exhibited a systematic decrease in crack propagation speed for peel angles smaller than ≈ 35 °. Dots represent detachments without visible sliding.

This result differs slightly from previous models for prestrained tape [25, 26, 27, 28, 29, 13], which we discuss in more detail in the Supplemental Material.

Strikingly, a similar, yet not identical result is obtained when the effect of frictional sliding (leading to pre-strain) is considered ([5, 18] and see Supplemental Material):

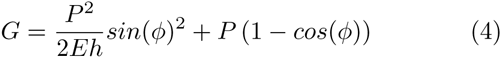

Equations 3 and 4 indicate that *identical* pre-tension at the peel front can lead to *different* peel strength if peeling is associated with interfacial slippage. This discrepancy is solely based on the fact that the peeled unit length refers to *unstretched* tape in the tape-sliding model, but to *stretched* tape in the pre-strain model (see Supplemental Material).

The difference between eqs. 3 and 4 is governed by a dimensionless parameter, *ζ* = *Eh/G,* which may be interpreted as the ratio of the elastic and adhesive work during peeling (see Supplemental Material). The two models are increasingly similar for large values of *ζ*, as both approach the inextensible tape model (eq. 2) as *ζ → ∞.* However, even moderately large values of *ζ* can lead to effectively inextensible behaviour. This can be illustrated with a simplified version of eq. 3, which can be found by assuming that the change in surface energy due to the additional peeled length arising from pre-tension is negligible (see Supplemental Material):

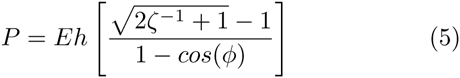

Equation 5 can be used for peeling without interfacial slippage, and is identical to the result given by refs. [25, 26, 27, 28, 29, 13] for *F*_0_ = *Fcos(ϕ).* Though incorrect, eq. 5 sets a conservative limit, and is reasonably close to the exact solution for large peel angles and *ζ >* 1 (see Supplemental Material), which is likely the case for most technical tapes and biological adhesives. The ratio of this force to the critical peel force for an inextensible tape is independent of the peel angle, and solely determined by *ζ*, i.e. adhesion tends to infinity as *ϕ →* 0. For a thin and soft tape with *G* = 100mNm^−1^, *h* =100*μ*m, and *E* =1 MPa, *ζ* = 10^3^, and the prediction of eq. 5 is within 0.005% of the inextensible tape model (eq. 2, see fig. 3). Even for very soft and thin structures, such as stick insect pads, the agreement is within 10% (see fig. 3). In practice, however, the yield strength of the tape may limit the force-enhancing effect of pre-tension considerably, and for elastomers, deformations may be sufficiently large to invalidate the assumption of linear elasticity. Models for large deformations can be found in [29, 18].

**Figure 3:**
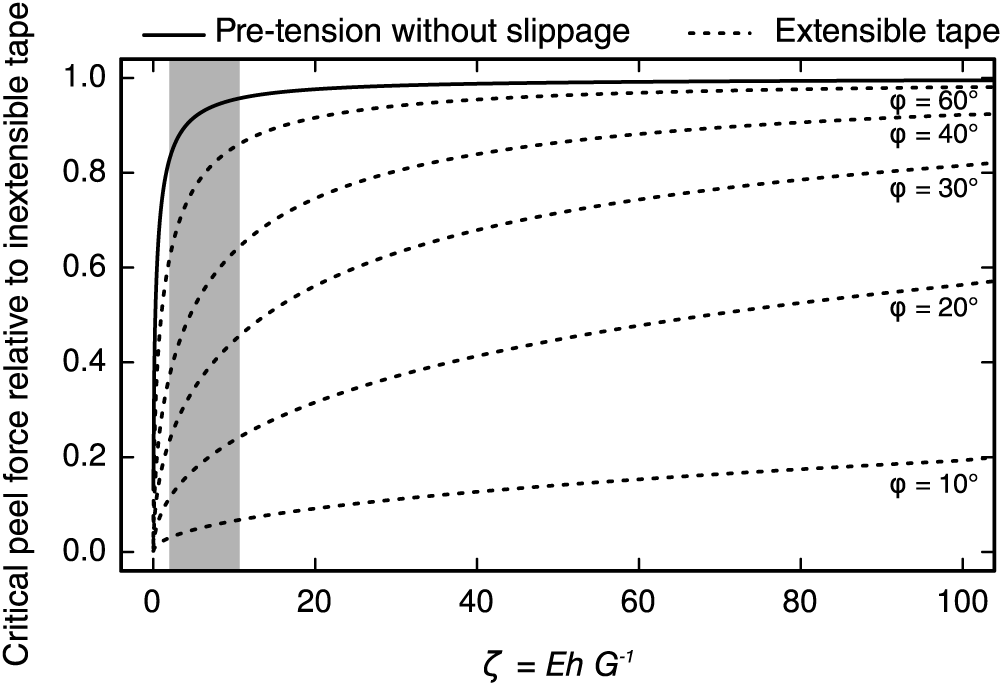
If adhesive tapes are pre-stretched with a force *F*_0_ = *Fcos(ϕ)* and peel without partial sliding, the critical peel force is an approximately constant fraction of the force required to peel an inextensible tape with identical critical energy release rate, independent of peel angle. Even for moderately large ratios ζ = *EhG^−^*^1^ as found in biological adhesive pads (grey box, see Supplemental Materials), the peel strength is close to that of an inextensible tape. As a comparison, dashed lines show the critical peel force for an extensible tape relative to the inextensible tape equation, for different peel angles.

Partial sliding during peeling will decrease the peel force in comparison to a tape with identical pre-strain at the peel front, but peeled without partial sliding. However, even with partial sliding, the peeling behaviour is similar to that of inextensible tape if *ζ* > 100 (see Supplemental Material). While the critical peel force for peeling with partial sliding is unbound as the peel angle approaches zero, the adhesive force per unit tape width, *Psin(ϕ),* remains finite and approaches 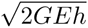. As *ϕ* → 0, an increasing fraction of the peel force acts in shear and is lost to sliding [18]. Hence, the critical energy required for crack propagation must be supplied by normal stresses, so that the *adhesive force* of peeling with sliding is approximately the same as the *total peel force* for an extensible adhesive tape without sliding (see eq. 1). When peeled at 0 ^°^, insufficient energy would be left to drive the crack through the interface, and instead the pads would merely slide [18].

The enhancing effect of pad pre-tension, both with and without interfacial slippage, readily justifies the use of the inextensible tape model even for small peel angles (fig. 3). The effect of pre-tension may also explain why a fit of the extensible tape model (eq. 1) to data from tree frogs required unrealistically large values for *E* [7]: Due to the pre-tension-induced changes in the peeling energy balance, even soft and thin pads can behave like an inextensible tape (fig. 3). This considerable increase in apparent stiffness during low-angle peeling is likely of major importance not only for the shear-sensitivity of smooth pads in stick insects, but for dynamic biological adhesives in general.

### Increase of the critical energy release rate via pad sliding

Tape pre-tension can result in a significant increase in the critical peel force in comparison to peeling of an unstretched pad, but it remains unclear whether it can fully account for shear-sensitive adhesion as observed in geckos, tree frogs, and stick insects. Notably, pre-tension of an adhesive tape prior to peeling can lead to a critical detachment angle [13], which seems consistent with observations on gecko adhesion [19, 6], and our data on stick insects. However, while all the modified peel models, including those for pre-strain and partial sliding (eqs. 3 and 4) predict forces *smaller* than for inextensible tape (see Supplemental Material), the adhesion forces we measured at low peel angles strongly *exceeded* this prediction (see fig. 1). This discrepancy is far from trivial: the adhesion forces predicted by the inextensible tape model (eq. 2) scale with the square root of the applied friction force for friction forces much larger than *Gw* [9]. However, we found an approximately linear relationship, i. e. the observed attachment forces (for large friction forces) exceeded the inextensible tape prediction by a factor approximately proportional to the square-root of the applied friction. In addition, a critical detachment angle does not occur if pad pre-stretch is based on sliding. Clearly, the shear-sensitivity of adhesion and the apparent critical detachment angle cannot be explained by any of the simple peeling models accounting for pre-tension and/or partial sliding. How then do sliding pads achieve forces much higher than the prediction of the inextensible tape model? Our findings on insects, and earlier results on geckos [6] can only be reconciled with peeling theory if the critical energy release rate *G* increases with the applied friction force (see eq. 2).

In fracture mechanics, *G* is often modelled as a function of ‘mode-mixity’, i. e. the extent to which interfacial failure occurs *via* shear versus tensile stresses (e.g. [32, 33]). For tape peeling, however, the mode-mixity dependence of *G* may largely arise from frictional ‘dissipation’ [18, 23], and is thus unlikely to provide an explanation (see above). Instead, the ‘true’ tensile strength of the interface must increase. Kendall [25] observed that crack-healing at low angles was accompanied by a significant increase of the critical energy release rate measured for *receding* cracks. Kendall suggested that this ‘surface activation’ may be partly explained by triboelectric charging, and indeed, sliding during tape peeling can lead to significant charges at the interface [34]. In order to test whether triboelectric charging can explain the observed increase of *G* for smaller peel angles, we repeated our experiments on grounded glass coverslips coated with conducting indium tin oxide. The relationship between friction and adhesion was virtually identical (see Supplemental Material). We therefore conclude that even if present, surface charging did not lead to a significant increase of *G*.

Adhesion depends on the ability of the interface to sustain stress. Insect adhesive pads are covered with a thin film of a secretion which acts as a separation layer, allowing to minimise viscoelastic losses during rapid detachments by providing a highly mobile interface through which a crack can easily propagate [10]. This effect, akin to slippage, reduces the critical stress concentration required for crack propagation, so that detachment forces remain small during voluntary detachment (see [35, 36, 37] for examples on synthetic adhesives). Pad sliding is accompanied by a loss of pad secretion at the pad’s trailing edge, which can lead to a significant increase in shear stress [38, 39, 40, 41]. A higher shear stress in a soft material implies an increase in adhesion hysteresis [42, 43], providing direct evidence for an increase in *G* upon reduction of the secretion film thickness (see also [44]). An increase in *G* as a result of sliding is also implied by the observation that crack propagation speed strongly decreased when pads slid during low-angle peeling, despite higher or equivalent normal stresses (see fig. 2B, and [45]). As the rate dependence of friction and bulk dissipation can differ considerably, even a minor increase in interfacial friction can change the adhesive force substantially [45]. An increase in *G* triggered by sliding may also be plausible for gecko setae [24], which have been shown to leave phospholipid footprints behind [46]; these could fulfil a similar function as the pad secretion in arthropods. Clearly, further research is required to investigate the role of interfacial mobility in biological adhesives.

Interface strengthening *via* sliding has at least two biologically relevant advantages over a typical peeling situation. First, as the onset of sliding depends on the pad’s contact area, not only friction but also adhesion forces may scale with pad area, which is increasingly difficult to achieve for larger animals [9]. Area scaling of adhesion is consistent with the observed, approximately linear relationship between friction and adhesion, and may be mediated by the pre-stretching of the pad, leading to a more uniform stress distribution across the pad contact zone. Second, the increase in *G* with friction force effectively expands the range of peel angles for which strong attachment is possible, but adhesive strength vanishes quickly when peel angles are larger than 30 ^°^, allowing a rapid switch (by a minimal change of the force direction) between strong attachment and effortless detachment during locomotion [6, 13].

## Conclusion

We have shown that the shear-sensitive adhesion in insects is consistent with classic peeling theory if friction forces are small, but a linear relationship between friction and adhesion occurs when friction forces are large. This coupling between adhesion and friction leads to a sharp increase of adhesion at peel angles smaller than 30^°^, which may result from two effects of sliding: First, partial sliding during detachment can give rise to considerable pre-tension, so that the pads have an increased apparent stiffness. Second, the thin films formed by the pad secretion result in a coupling of interfacial and bulk properties: pad sliding reduces the thickness of the fluid layer in the contact zone, and the interface now has a lower mobility, so that slippage is reduced, and stresses need to rise to higher levels to drive detachment. Larger stress levels increase the deformed volume of the adhesive pad, thereby increasing bulk dissipation within the adhesive pad cuticle [22]. As a consequence, peel forces exceed the predictions for an inextensible tape with constant critical energy release rate *G*. In combination, these effects may explain the sharp increase of adhesion with decreasing peel angle, and the approximately linear relationship between adhesion and friction observed in dynamic biological adhesives, allowing climbing animals to switch rapidly between attachment and detachment.

Our results demonstrate that the impressive controllability of biological adhesives does not solely arise from the pads’ structural anisotropy and direction-dependence, but is directly linked to processes at the interface. This suggests a promising new route for the development of bioinspired adhesives with simple morphology, but high controllability. Most technical adhesives are polymers, whose interfacial properties can be fine-tuned on a molecular scale. The extensive theoretical and experimental toolbox available to study and model the adhesion of polymers [47, 45] should allow to create technical adhesives with similar interfacial properties, replicating some of the most desirable features of biological adhesives.

## Materials and methods

Attachment performance of single pads of live Indian stick insects *(Carausius morosus,* Sinety, 1901; mass=618 ± 101mg, mean ± s.d., n=11) was measured with a custom-made 2D-force transducer set-up described in detail in Drechsler and Federle [39] (see fig. 4 for a schematic of the set-up). The pads were mounted using the method described in Labonte and Federle [8]. During the force measurements, the contact area of the pads was recorded with a Redlake PCI 1000 B/W high-speed camera (Redlake MASD LLC, San Diego, CA, USA), mounted on a coaxially illuminated stereo-microscope (Leica MZ16, Leica Microsystems GmbH, Wetzlar, Germany). All measurements were conducted at 22-24° C and 40-50% humidity, and with clean glass or indium tin oxide (ITO) coated coverslips purchased from Diamond Coatings Ltd. (Halesowen, UK). The ITO coverslips had a resistance of 15-30Ω, measured with electrodes attached on opposite sides of the 18 × 18 mm coverslips (Fluke 27 multimeter, RS Components Ltd, Corby, UK); the coverslips were grounded during the force measurements.

**Figure 4:**
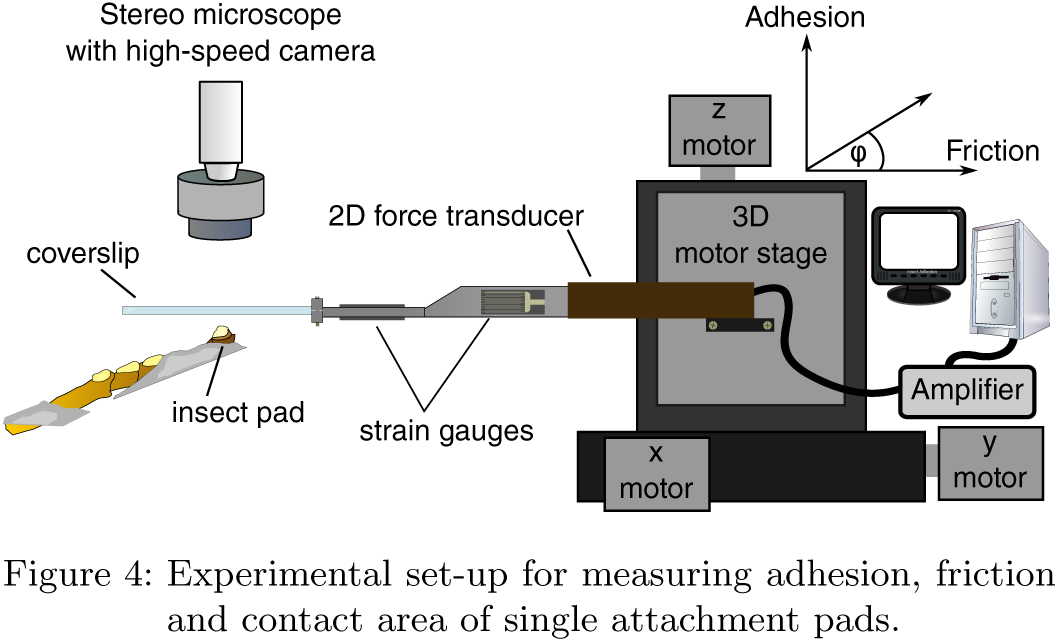
Experimental set-up for measuring adhesion, friction and contact area of single attachment pads.

### Measurement protocol

Peak adhesion of stick insect arolia was measured under two different conditions for all specimens: (i) by retracting the coverslips with constant speed and different constant ‘retraction angles’, altered by adjusting the movement velocity of each motor axis and (ii), by retracting the coverslips perpendicularly after a defined shear force was applied to the pads [8]. The order of the conditions was randomized, and each measurement was performed on a fresh position of the surface, in order to avoid a systematic influence of fluid accumulation and/or depletion [39, 40].

For the first measurement series, the surface was initially pressed onto the pads with a normal preload of 1 mN, corresponding to approximately 1/6^th^ of the body weight, controlled via a motorized 20 Hz force-feedback mechanism incorporated in a custom-made Labview control software (National Instruments, Austin, TX, USA). After a contact time of 5 s, the surface was retracted at a defined retraction angle (given by the motor trajectory), with a constant motor speed along the trajectory of 0.5 mm s^−1^.

Measurements were performed for nine retraction angles, ranging from 90 to 10 ° in steps of 10° (here, 90 ° corresponds to a perpendicular pull-off). For the second series of measurements, the surface was pressed onto the pads with a preload of 1mN for 5 s as before. Subsequently, the motorised force-feedback mechanism was used to apply a constant shear force for a period of 3 s, followed by a perpendicular pull-off at 0.5mms^−1^ (i.e. during detachment, the beam was only moved perpendicularly by the motors). Measurements were performed for eight different shear forces, ranging from 5% to 170% of the body weight (0.25, 0.5, 1, 2, 4, 6, 8 and 10mN).

Force-displacement data were recorded with an acquisition frequency of 20 Hz, and the pad contact area was filmed at 200 frames per second for shear-force feedback experiments, or at 100 frames per second for the measurements that involved detachment at defined retraction angles. The difference in framerate was owing to the limited memory of the camera and the longer times required to detach the pads at peeling angles <30 °.

For both types of measurements, peak adhesion and the friction forces at this peak were extracted from the force-time curves. From these data, we also calculated a ‘force’ peel angle, i.e. the arc tangent of the ratio of both forces. As the relationship between peel angle and adhesion did not differ between the two types of experiments, the data were pooled (repeated measures ANCOVA, *F*_1,192_ =2.39, p=0.12, n=11 for both types of measurements).

In order to measure the width *w*, area *A*, and perimeter *Γ* of the pad contact area, the video recordings were post-processed using Fiji [48]. Video recordings were filtered in order to remove flickering from the light source and subsequently converted into binary images using the fuzzy threshold algorithm described in ref. [49]. The binary images were despeckled using 2 × 2-5 × 5 pixels median filters and the resulting stacks were analysed with the native particle analysis routines implemented in ImageJ 1.48k.

From the processed videos, we also measured the speed of crack propagation *v_c_* [31]

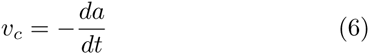

where *a* = *A/Γ* is the contact radius. *v_c_* changed during detachment, and we measured the ‘terminal’ speed of crack propagation by performing an ordinary least square regression of contact radius against time for the last 50 ms of detachment (i.e. for 5 and 10 data points for 100 and 200Hz recordings, respectively). From the video recordings, we also determined whether pads were sliding during detachment, which was clearly visible as a change of the pad position relative to features on the coverslips.

All statistical analysis was carried out with *R* v.3.0.2 [50].

